# YTHDF1 drives intestinal immune response against bacterial infection

**DOI:** 10.1101/2020.05.08.083840

**Authors:** Xin Zong, Xiao Xiao, Bin Shen, Qin Jiang, Hong Wang, Zeqing Lu, Fengqin Wang, Minliang Jing, Yizhen Wang

**Author notes:** Correspondence: 866 Yuhangtang Road, Hangzhou, Zhejiang Province 310058, People’s Republic of China.

## Abstract

Invasion of pathogenic bacteria is a serious threat to intestinal health. Recent emerging evidence has demonstrated that N6-methyladenosine (m^6^A) is closely associated with innate immunity; however, the underlying mechanism remains unclear. Herein, we aim to explore the function and mechanism of m^6^A modification in the regulation of innate immune responses against bacterial pathogens in the intestine. Ribo-seq and m^6^A-seq data have demonstrated that YTHDF1, an m^6^A reader, directs the translation of tumor necrosis factor receptor-associated factor 6 (TRAF6) mRNA to regulate immune responses via modulation of m^6^A methylation near stop codon. Furthermore, we have identified a unique mechanism that the interaction between YTHDF1 and the host factor DDX60 are critical in regulating intestinal immune response against bacterial infection by recognizing TRAF6 target transcripts. Additionally, our results provide novel insights as to why YTHDF1 could recognize its unique targets using the same domain as other YTHDF proteins. This work identifies YTHDF1 as a key driver of intestinal immune responses and provides an avenue for development of novel strategies to modulate intestinal immune response against bacterial infection.

## Introduction

Enterotoxigenic *Escherichia coli* (*ETEC*) causes diarrhea by colonizing the small intestine, leading to substantial mortality and morbidity in children and animals (Qadri, Akhtar et al., 2019). Intestinal epithelial cells (IECs) constitute an integral component of a highly regulated communication network that senses microbial and environmental stimuli, as well as endogenous danger signals (Peterson & Artis, 2014). Activation of an IEC-specific immune response induces the production of a myriad of cytokines, chemokines and acute phase proteins (Henderson, van Limbergen et al., 2011). It is well known that toll-like receptors (TLRs) and nuclear factor-κB (NF-κB) signaling play important roles in cytokine and chemokine production to induce the epithelial immune defense, mobilize immune effector cells, and activate adaptive immunity (Price, Shamardani et al., 2018). Tumor necrosis factor receptor-associated factor 6 (TRAF6), one of the key regulators mediating TLR and subsequent NF-κB signaling, is triggered by complex stimuli, including bacterial infection (Walsh, Lee et al., 2015). However, the mechanisms that regulate immune responses in the intestine remain largely unexplored.

Recently, increasing evidence has indicated that RNA modifications provide a powerful means to dynamically regulate gene expression (Zhao, Roundtree et al., 2018). Methylation of adenosine at the N6 position (m^6^A) is the most prevalent known posttranscriptional modification in mammalian mRNA (Roundtree, Evans et al., 2017). Similar to DNA methylation, m^6^A modification is reversibly catalyzed by a different set of enzymes; the modification is generated by a stable core catalytic complex (METTL3/METTL14/WTAP) and can be removed by demethylase FTO or ALKBH5. This m^6^A functions as a “reader” that directly binds to RNA in the cytoplasmic space (Zhang, Fu et al., 2019). To date, three major “reader” proteins, including those of the YTH domain family (YTHDF) 1, YTHDF2, and YTHDF3, have been demonstrated to recognize the m^6^A site through their YTH domain (Shi, Wei et al., 2019). Among these, YTHDF1 is believed to interact with translation initiation factors during the translation of m^6^A-modified mRNA (Wang, Zhao et al., 2015), whereas YTHDF2 promotes the degradation of its target transcripts partially via recruitment of the CCR4-NOT deadenylase complex (Du, Zhao et al., 2016). YTHDF3 cooperates with both YTHDF1 and YTHDF2 to regulate the translation and decay of methylated mRNAs (Shi, Wang et al., 2017). However, the role(s) of YTHDF proteins in IEC-specific immune responses and the mechanism underlying the triggering of YTHDF protein by stimuli to promote translation or degradation remain unknown. Function of RNA m^6^A modification as well as YTHDF proteins in intestinal immune responses is of interest; however, it has not been investigated in detail.

DEAD-box (DDX) proteins, which contain a conserved Asp-Glu-Ala-Asp motif, are the largest family of RNA helicases (Banroques, Cordin et al., 2011). DDX proteins interact with both rRNAs and mRNAs to regulate multiple biological functions, such as translation initiation, mRNA synthesis, and RNA splicing (Fu, Wu et al., 2016). The DEAD-box helicase DDX60 was identified from virus-infected human dendritic cells (DCs) (Miyashita, Oshiumi et al., 2011); it is involved in a subset of innate immune pathways across various cell types (Goubau, van der Veen et al., 2015). In this study, we revealed that YTHDF1 is critical for intestinal immune responses against bacterial infection both *in vitro* and *in vivo*. Apparently, YTHDF1 ablation resulted in substantially reduced immune responses via downregulation of TRAF6 transcript translation. METTL3-mediated m^6^A modifications, either within the 3**′**UTR region or the coding sequence (CDS), are involved in these events. We also showed that interaction with DDX60 is essential for YTHDF1 to recognize m^6^A on TRAF6 transcripts and to direct its protein synthesis. Collectively, our findings highlighted an important role of YTHDF1 in protection against bacterial infection and maintenance of intestinal immune homeostasis.

## Results

### YTHDF1 is critical for IECs to defense against bacterial infection

To evaluate the potential involvement of the YTHDF protein family in IECs upon bacterial infection, we first investigated cultured IPEC-J2 cells exposed to LPS and *ETEC*. We found that gene copy number of YTHDF1 was much higher than that of YTHDF2 or YTHDF3 after LPS stimulation or *ETEC* infection (Fig 1A), which indicated that YTHDF1 plays an important role in these events.

**Fig 1.**
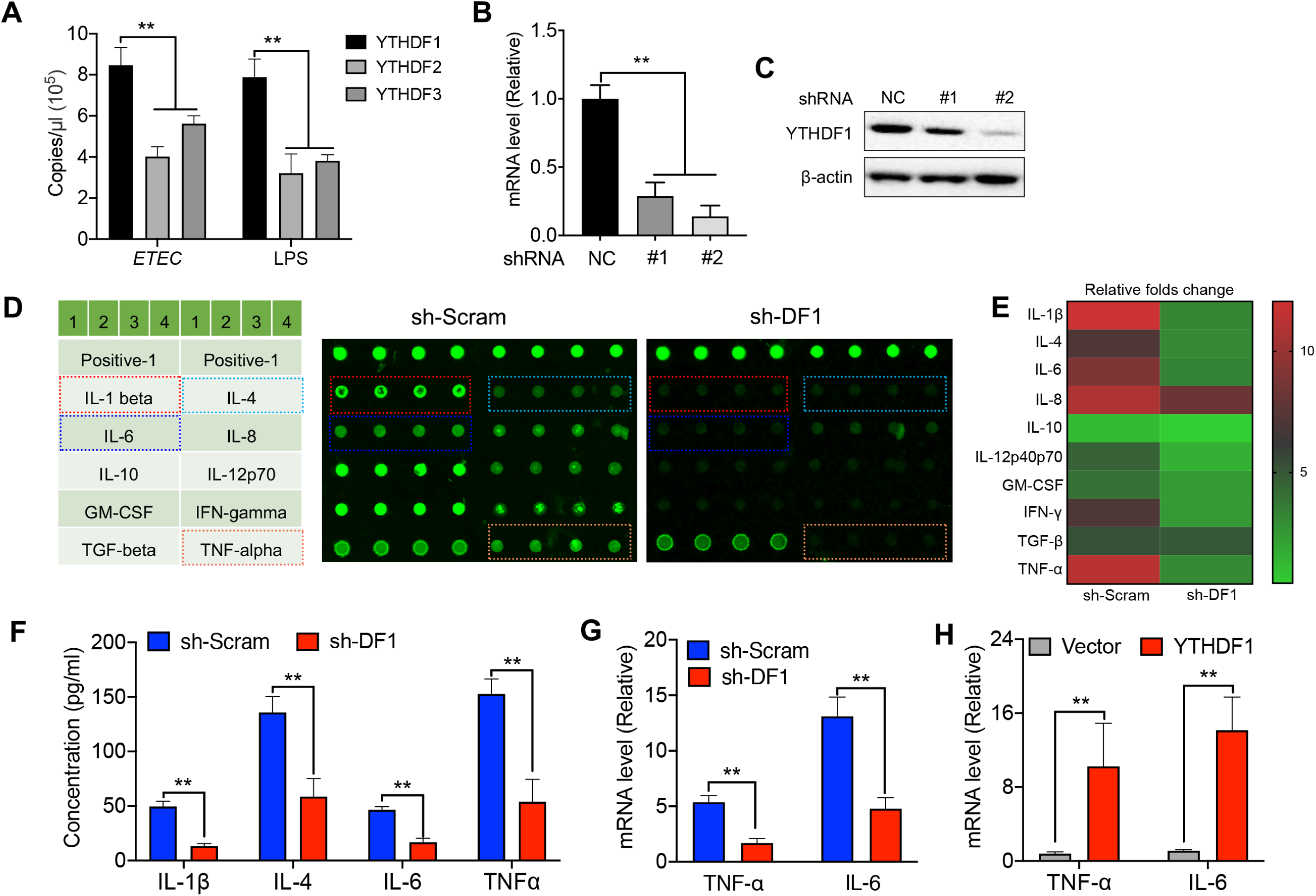
YTHDF1 mediated the bacterial immune response in IECs. (A) Gene copy numbers of YTHDF1, YTHDF2, and YTHDF3 after LPS stimulation or ETEC infection. (B-C) The diminished effects of an established YTHDF1 depletion in IPEC-J2 cells; qPCR analysis of mRNA level (B); immunoblot analysis of protein level (C). NC, cells transfected with negative control scramble shRNA; shYthdf1, cells transfected with YTHDF1 shRNA [one of two shRNA constructs (1, 2)]. (D) Laser scanning map of the cytokine microarray for LPS-treated indicated cells. Each cytokine was arrayed in quadruplicates (left table indicates cytokine map). Dots on the top represent positive controls. (E) The relative fold change by signal intensity from (D). The results are relative to those of sh-Scramble group. (F-G) ETEC-infected IPEC-J2 cells with 3 h treatment, cytokines in supernatants (F), and mRNA levels (G) were determined. (H) TNF-α and IL-6 mRNA levels in YTHDF1-depleted IPEC-J2 cells added back with the YTHDF1 plasmids. The qPCR results are presented relative to those of GAPDH. The data are expressed as the mean ± s.e.m.; ^*^*P* < 0.05, ^**^*P* < 0.01, n = 3 biological replicates.

To determine the role of YTHDF1 in IECs’ defense against bacterial infection, we silenced YTHDF1 expression in IPEC-J2 cells by lentiviral vector-delivered expression of short hairpin RNAs (Fig 1B and C). Remarkably, cytokine multiplex assays showed that LPS-induced cytokine production, including that of IL-1β, IL-6, and TNF-α, was substantially decreased under YTHDF1-deficient condition (Fig 1D and E). These results were verified by ELISA and qPCR (Fig 1F and G). Similarly, *ETEC*-induced expression of inflammatory cytokines, such as TNF-α and IL-6, were downregulated owing to YTHDF1 deficiency (Fig EV1A and B). This phenotype was also validated in the human Caco-2 cells (Fig EV1C). In addition, we conducted rescue experiments in YTHDF1-depleted IPEC-J2 cells to further confirm that the decreased cytokine signature was specific to YTHDF1 knockdown. As expected, we found that the reintroduction of YTHDF1, not the vector control, enhanced cytokine expression upon infection in YTHDF1-depleted cells (Fig 1H). Furthermore, we observed that depletion of YTHDF1 suppressed activation of the transcription factor NF-κB (p65) during LPS and *ETEC* stimulation, suggesting that YTHDF1 functions upstream of p65 (Fig EV2A and B). These data indicate that YTHDF1 is important for the immune response to combat infection in IECs, possibly via regulation of NF-κB signaling.

### YTHDF1 regulates IECs immune responses by directing translation of TRAF6

YTHDF1 primarily functions in ribosomal loading of m^6^A-modified mRNAs to facilitate translation initiation (Wang et al., 2015). To explore the downstream targets of YTHDF1, we employed ribosome profiling to compare the translation efficiency profile in LPS-stimulated IPEC-J2 cells, either in the presence or absence of YTHDF1. As expected, compared with the scramble control cells, we found a notable decrease in translation efficiency, particularly for m^6^A-modified transcripts, in YTHDF1-knockdown IPEC-J2 cells (Fig 2A and B). A total of 2030 differentially translated mRNAs (947 upregulated and 1083 downregulated genes) were selected for gene KEGG pathway enrichment analysis (Fig 2C). We found that among the downregulated genes the top enriched pathways were involved in the following: “TNF signaling pathway,” “Salmonella infection,” and “NF-κB signaling pathway” (Fig EV3). Because previous results indicated that YTHDF1 might play a role in NF-κB signaling and is upstream of p65, we focused on the NF-κB signaling pathway among the group of downregulated genes. We observed that the translation efficiency of TRAF6 mRNA was significantly decreased (Fig 2D), whereas that of IκBα was merely affected (Fig EV4A). To directly demonstrate the subsequent translational outcomes of TRAF6, we performed a firefly luciferase (Fluc) reporter assay in IPEC-J2 cells in either the absence or presence of YTHDF1. Consistent with ribosome profiling, a significant decrease in TRAF6 translation was YTHDF1 is knockdown (Fig 2E). Furthermore, we analyzed the protein synthesis of TRAF6 with or without YTHDF1. Remarkably, TRAF6 synthesis was substantially reduced in the absence of YTHDF1 (Fig EV4B). Comparable levels of TRAF6 mRNA indicated that such reduced TRAF6 synthesis resulted from translational deficiency (Fig EV4B). Similar observations were also made following *ETEC* infection in human Caco-2 cells (Fig EV4C and D). Because TRAF6 is a key regulator in mediating TLR4 and subsequent signaling, we also analyzed the effect of YTHDF1 depletion on TLR4 signaling. Expectedly, downstream targets of TRAF6 such as IKK, JNK, and MAPK were suppressed, whereas those upstream of TRAF6 remained unchanged (Fig EV5). Cumulatively, we speculated that TRAF6 mRNA is one of the key targets of YTHDF1 in governing immune responses.

**Fig 2.**
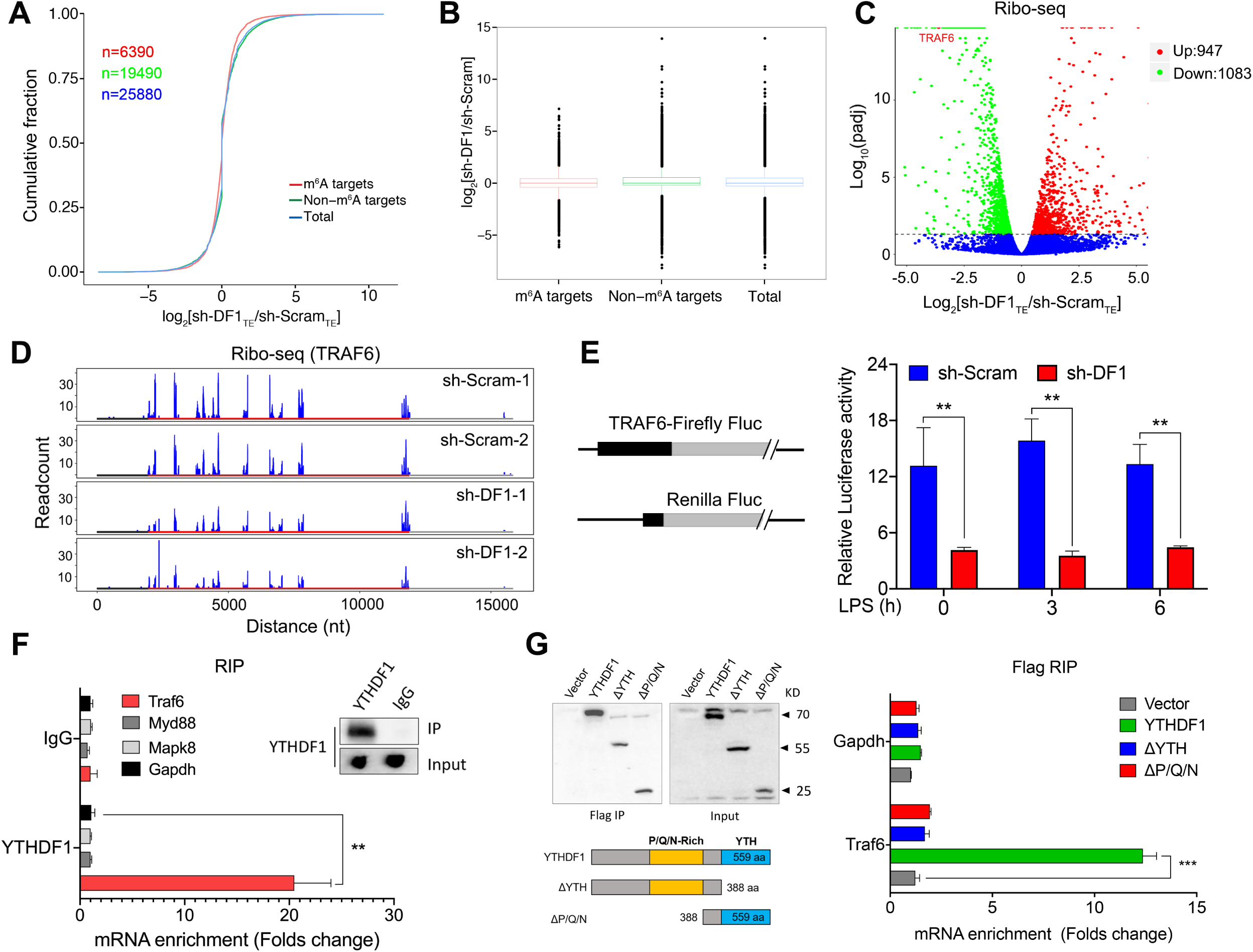
YTHDF1 maintained TRAF6 synthesis by directing its translation initiation. (A) The cumulative distribution of the fold change showing the translation efficiency between LPS-treated sh-Scramble (sh-ScramTE) and shDF1 (sh-DF1TE) IPEC-J2 cells for m^6^A targets (red), non-m^6^A targets (green), and total (blue) mRNA. *P*-values were calculated using a two-sided Kolmogorov–Smirnov test; n = 2 independent biological replicates. (B) Box plot depicts the cumulative distribution of (A). Boxplot elements: center line, median; box limits, upper and lower quartiles; whiskers, 1%–99%. (C) Volcano plots of genes with differential translation efficiency from the same samples as in (A). The *P*-values were calculated with a two-sided likelihood ratio test and adjusted by the Benjamini–Hochberg method. (D) Genome-wide ribosome profiling of ribosome-protected mRNA fragments for TRAF6 in LPS-treated IPEC-J2 cells with or without YTHDF1 (sh-Scram or sh-DF1, respectively). Scales represent genomic RNA size and structure. (E) Luciferase activity of TRAF6 in IPEC-J2 cells after transfection with the indicated plasmids. Luciferase activity is presented relative to Renilla luciferase activity. (F) The interaction between YTHDF1 and TRAF6 transcripts was measured by RNA immunoprecipitation (RIP)-qPCR and normalized to the input levels. Immunoblot of YTHDF1 (top right) showed that YTHDF1 antibody was immunoprecipitated successfully. (G) RIP-qPCR analyzed TRAF6 and GAPDH abundance immunoprecipitated by Flag antibody from the IPEC-J2 cells overexpressing YTHDF1, ΔYTH, and ΔP/Q/N plasmid. The data are expressed as the mean ± s.e.m.; ^**^*P* < 0.01, ^***^*P* < 0.001, n = 3 biological replicates.

RNA binding activity is necessary for YTHDF1 to regulate translation. Thus, we performed the RNA immunoprecipitation (RIP)-qPCR to confirm the presence of this specific binding mechanism between YTHDF1 and TRAF6 mRNA. As expected, in contrast to the control immunoglobulin G (IgG), we detected abundant TRAF6 mRNA signal, but not for MyD88 and MAPK8, upon immunoprecipitation with YTHDF1 antibody (Fig 2F). These results indicated that the inactivation of MAPK signaling was due to suppressed TRAF6 expression, and was not a direct result of a lack of YTHDF1. Next, we constructed a YTHDF1-truncated plasmid and found that deletion of either the YTH domain (ΔYTH) or Pro/Gln/Asn-rich domain (ΔP/Q/N) of YTHDF1 abrogated its binding to TRAF6 mRNA (Fig 2G). Furthermore, all YTHDF1-truncated plasmid failed to rescue the decreased cytokine expression in YTHDF1-silenced IPEC-J2 cells (Fig EV6). These data suggest that YTHDF1 binds to TRAF6 mRNAs to initiate their translation and thereby regulate the expression of cytokines, and both the YTH and P/Q/N-rich domains of YTHDF1 are indispensable for effective YTHDF1-RNA interactions.

### YTHDF1-mediated translation of TRAF6 requires m^6^A methylation

As aforementioned, YTHDF1 is a “reader” of m^6^A. To test whether altered YTHDF1-sensitive gene expression is associated with m^6^A modifications, we first compared m^6^A abundance in scramble controls and YTHDF1-depleted IPEC-J2 cells with LPS stimulation. In contrast to the globally increased m^6^A levels observed in the control cells upon LPS treatment, cells lacking YTHDF1 surprisingly showed no elevated levels of methylation (Fig EV7A), suggesting the involvement of mRNA methylation in YTHDF1-sensitive gene expression. Because YTHDF1 is not a methyltransferase, we determined the levels of methyltransferase and m^6^A demethylase in cells exposed to LPS, which revealed that loss of YTHDF1 led to decreased levels of METTL3 but increased levels of FTO (Fig EV7B). Next, we mapped the m^6^A methylomes in IPEC-J2 cells before and after infection using m^6^A sequencing (m^6^A-Seq). We searched for consensus motifs and identified a GGACU sequence (Fig 3A), which was consistent with a previous report (Zhou, Wan et al., 2018). Among 240 substantially different methylated transcripts, 141 (including TRAF6) were upregulated, whereas the other 89 were downregulated (Fig 3B). In addition, similar pattern of m^6^A distribution was observed in control and infected cells, regardless of whether the transcripts were upregulated or downregulated, with a majority of m^6^A modifications being enriched near a stop codon and within the 3**′**UTR (Fig EV8A). Notably, we identified three statistically significant m^6^A peaks in the TRAF6 transcripts, with two in the CDS vicinity of the stop codon and one representing a unique peak within the 3**′**UTR. All these peaks were substantially increased post-infection, suggesting functional relevance (Fig 3C). As expected, methylation levels of m^6^A, as indicated by MyD88 mRNA, showed no difference between control and infected cells (Fig EV8B). These observations suggest that m^6^A modification on TRAF6 mRNAs is enhanced upon bacterial infection and is involved in its translation.

**Fig 3.**
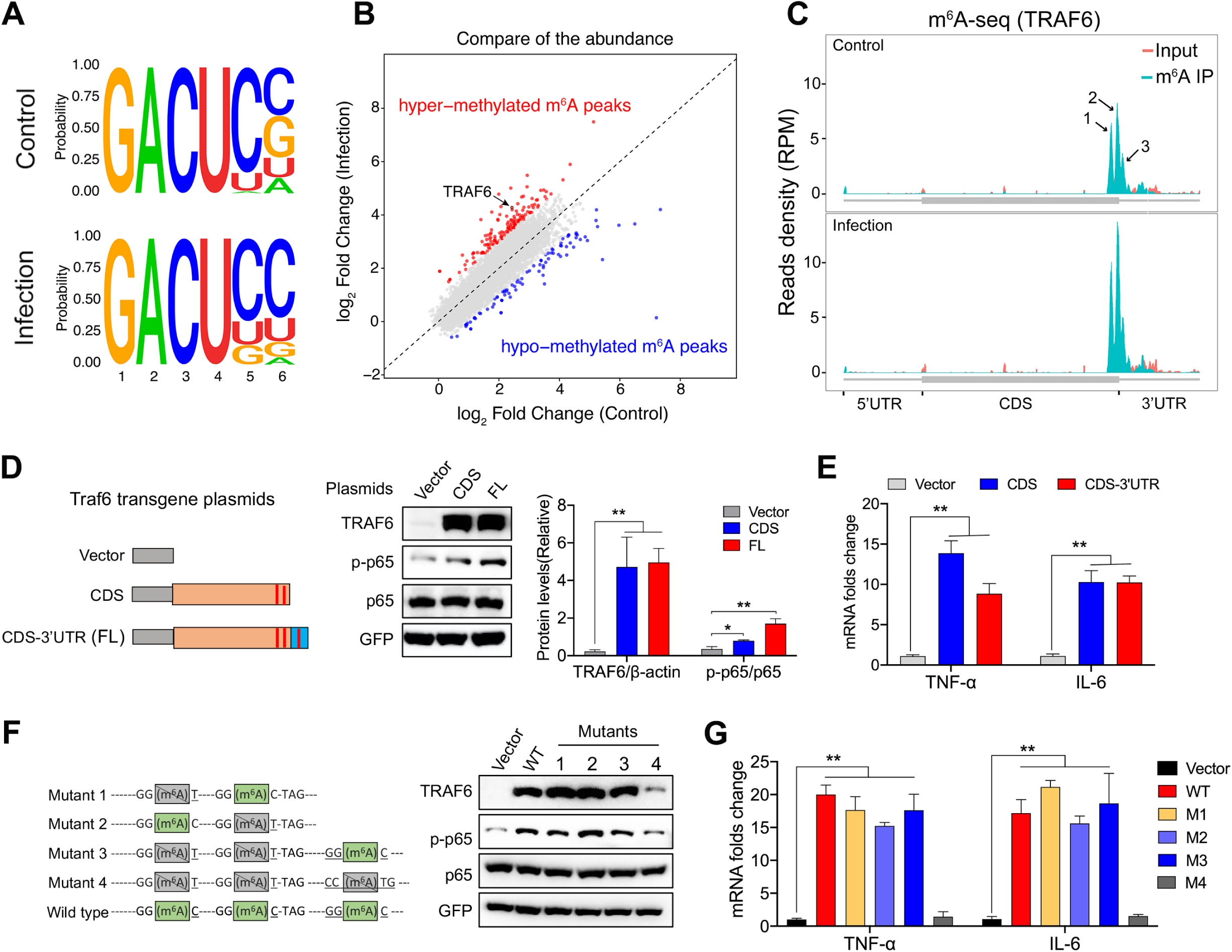
Translation of TRAF6 was dependent on m^6^A methylation. (A) Sequence motifs within m^6^A peaks identified by using Homer software. (B) Analysis of differential m^6^A-modified gene in either uninfected IPEC-J2 cells or those infected with *ETEC*. (C) RNA-Seq of input RNA and m^6^A immunoprecipitated (IP) RNA in replicates for the TRAF6 transcript. The m^6^A motif sequences that correspond to an immunoprecipitate-enriched region are marked in green. (D) Rescue assay of specific TRAF6 plasmid in LPS-treated TRAF6-silenced IPEC-J2 cells. The left panel shows strategies of various TRAF6 plasmid construction. All plasmids were cloned into the FLAG-pcDNA3.1 vector. The right panel shows relative protein levels quantified by densitometry and normalized to the level of GFP. (E) TNF-α and IL-6 mRNA levels in TRAF6-depleted IPEC-J2 cells added back to the specific TRAF6 plasmid from (D). (F) Immunoblot analysis of FLAG-TRAF6, p65, and p-p65 in TRAF6-depleted IPEC-J2 cells transfected with specific TRAF6 plasmids and treated with 50 μg/ml LPS for 6 h. The left panel shows the mutation strategy of the m^6^A sites in TRAF6 mRNA. All plasmids were cloned into the FLAG-pcDNA3.1 vector. (G) TNF-α and IL-6 mRNA level in TRAF6-depleted IPEC-J2 cells were added back to the various TRAF6 plasmid from (F). The qPCR results are presented relative to those of GAPDH. The data are expressed as the mean ± s.e.m.; ^**^*P* < 0.01, n.s., not significant, n = 3 biological replicates.

To further analyze the function of m^6^A modification on TRAF6 mRNA, we created FLAG-tagged expression constructs containing the TRAF6 CDS with or without the 3**′**UTR (CDS-3**′**UTR) (Fig 3D). After transfection into TRAF6-silenced IPEC-J2 cells, we found that protein expression of both TRAF6 constructs was significantly increased in contrast to vector following ETEC infection (Fig 3D), suggesting that the 3**′**UTR or rather the m^6^A within the 3**′**UTR was dispensable for YTHDF1–TRAF6 mRNA regulation. The enhanced expression of both TNF-α and IL-6 induced by the forced expression of FLAG-tagged TRAF6 further confirmed this conclusion (Fig 3E). Together, these data indicate that TRAF6 protein synthesis depends on both YTHDF1 and m^6^A modification in the CDS region of its mRNA.

Next, we introduced a mutation in the m6A sites (corresponding to the conserved m6A motif) on TRAF6 mRNA (Fig 3F) in the FLAG-TRAF6 plasmids. These constructs were then transfected into TRAF6-silenced IPEC-J2 cells. After ETEC infection, we found that the TRAF6 protein levels were remarkably increased in cells transfected with mut1-3, but not mut4 (all three m6A sites mutated) (Fig 3F). Consistently, in addition to the mut4 plasmid, the other three mutation plasmids had similar effects as wild-type TRAF6 plasmids upon inducing the expression of cytokines (Fig 3G). In short, in a bacterial immune response, m^6^A near the stop codon, either within the CDS or 3′UTR, is capable of mediating the mRNA translation of TRAF6.

### DDX60 is a coregulator for YTHDF1 regulating TRAF6 synthesize

To gain further insights into the mechanism of YTHDF1–TRAF6 mRNA action, we hypothesized that the presence of a coregulator is required for facilitating YTHDF1 to accurately recognize its targets and such an effector molecule is sensitive to the states of both YTHDF1 and m^6^A. As an alternative approach, we purified YTHDF1-interacted proteins using YTHDF1 antibody obtained from LPS-stimulated IPEC-J2 cells. We employed tandem mass spectrometry analysis (MS/MS) to identify and quantify the associated protein components. In total, 466 proteins were enriched by YTHDF1, with the top candidate being DDX60 (Fig 4A). Next, we directly confirmed the interaction between YTHDF1 and DDX60 by immunoprecipitation and identified an endogenous interaction between YTHDF1 and DDX60 in LPS-stimulated cells. Moreover, the interaction between YTHDF1 and DDX60 persisted even after RNA degradation by RNase (Fig 4B and C).

**Fig 4.**
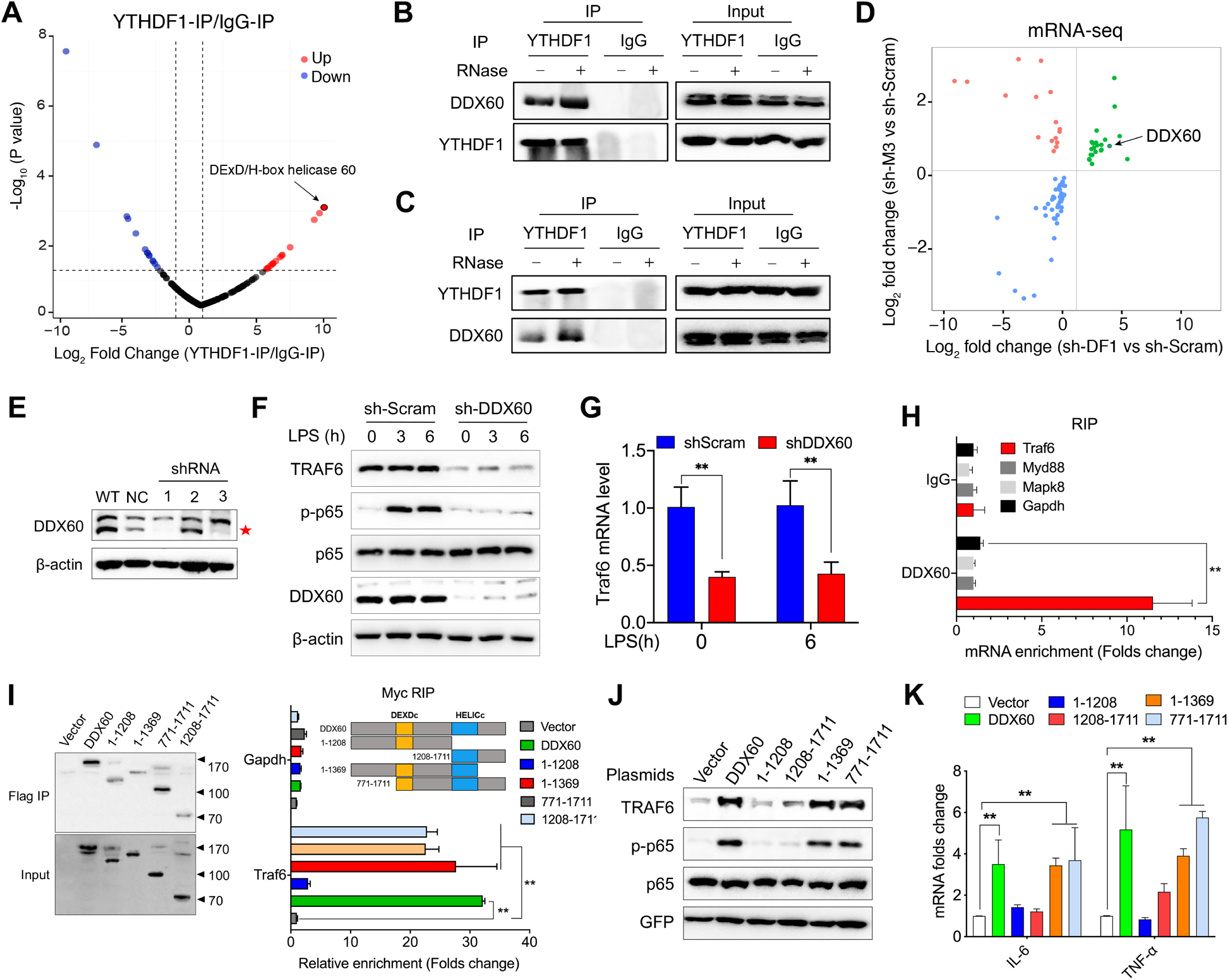
DDX60 involved in TRAF6 protein synthesis. (A) Volcano plot of the Log10 (*P*-value) versus Log2 fold change (YTHDF1-IP/IgG-IP) of MS-identified proteins from two biological replicates. Intensities were obtained from a MaxQuant database search. (B-C) Immunoprecipitation was performed with antibody to YTHDF1 (B) or DDX60 (B) and control IgG. After immunoprecipitation, samples were washed and incubated with RNase as indicated. The samples were analyzed by immunoblot assay using anti-YTHDF1 and anti-DDX60 antibodies. (D) Significantly regulated genes (*P* < 0.05) from overlapped YTHDF1 knockdown and METTL3 knockdown IPEC-J2 cells are presented in a scatter plot showing the genes that were commonly upregulated or downregulated following either YTHDF1 or METTL3 silencing (highlighted in green and blue, respectively). (E) Immunoblot analysis of DDX60 in IPEC-J2 cells transfected with negative control shRNA (NC) or DDX60-specific shRNA (one of three shRNA constructs: 1, 2, or 3). (F) Immunoblot analysis of TRAF6 and its downstream targets in indicated cells after LPS stimulation. The right panel shows the relative protein levels of TRAF6 quantified by densitometry and normalized to the level of β-actin. (E) Interaction between DDX60 and TRAF6 transcripts was measured by RIP-qPCR and normalized to the input levels. Immunoblot of DDX60 (top right) showed that DDX60 antibody was immunoprecipitated successfully. (F) The fold enrichment of TRAF6 and GAPDH mRNA as determined by RIP-qPCR in IPEC-J2 cells overexpressed the indicated plasmids. Schematic structure of Myc-DDX60 and the truncation strategy is shown above. (G) The shDDX60 cells added back to the indicated plasmids upon LPS stimulation. The protein level of TRAF6, p65, and p-p65 were determined by immunoblot. The right panel shows the relative protein levels quantified by densitometry and normalized to the quantity of β-actin. The qPCR results are presented in relation to those of GAPDH. The data are expressed as the mean ± s.e.m.; ^**^*P* < 0.01, ^***^*P* < 0.001, n.s., not significant, n = 3 biological replicates.

This coregulator is sensitive to not only YTHDF1 but also m^6^A. To further confirm the possibility that DDX60 is the coregulator, the transcriptome was analyzed by mRNA sequencing (mRNA-seq) using RNA isolated from scramble control, YTHDF1-knockdown, and METTL3-knockdown IPEC-J2 cells. METTL3 is a core methyltransferase of m^6^A and determines the global level of mRNA m^6^A modification (Liu, Yue et al., 2014). DDX60 was analyzed in the DEGs sensitive to both YTHDF1 and METTL3, which revealed that DDX60 may act as a coregulator in YTHDF1–m^6^A interaction(s) (Fig 4D). Furthermore, loss of YTHDF1 led to increased DDX60 protein level, which was consistent with the mRNA-seq data (Fig EV9A).

Based on the aforementioned findings, we speculated that DDX60 interacts with YTHDF1 and is involved in YTHDF1–TRAF6 regulation. To test this hypothesis, we created a DDX60-knockdown IPEC-J2 cell line (Fig 4E). Upon LPS stimulation, the DDX60 knockdown suppressed TRAF6 synthesis and blocked activation of the NF-κB signaling pathway as compared with the scramble control (Fig 4F). We observed a similar phenotype in human Caco-2 cells and ETEC*-*infected IPEC-J2 cells (Fig EV9B and C). Moreover, decreased production and gene expression of TNF-α and IL-6 confirmed these observations (Appendix Fig EV9D-F). Surprisingly, in contrast to the YTHDF1 knockdown, the transcription of TRAF6 was also decreased in cells lacking DDX60 (Fig 4G). These observations indicate that DDX60 may act as a coregulator via stabilization and transportation of TRAF6 mRNAs to ensure that YTHDF1 accurately recognizes TRAF6 mRNA.

### DDX60 facilitates of YTHDF1 to accurately recognize TRAF6 mRNA

Based on the working model of DDX60, we further tested whether DDX60 interacts with TRAF6 transcripts. Similar to YTHDF1, the abundance of TRAF6 transcripts but not those of MyD88 and MAPK8 was enriched by DDX60 (Fig 4H). Next, we constructed additional plasmids expressing various truncated fragments of DDX60 (Fig 4I). RIP-qPCR revealed that DDX60 lacking the HELICc domain could no longer enrich TRAF6 transcripts (Fig 4I), indicating that this domain is involved in its RNA binding activity. Furthermore, neither the DEADc nor HELICc domains alone could rescue the suppressed expression of TRAF6 or the activation of p65 caused by deletion of DDX60 (Fig 4J). Further functional studies showed a noticeable increase in cytokine expression in DDX60-depleted cells that were transfected with truncated DDX60 plasmids containing DEADc and HELICc domains, but not in cells transfected with plasmids containing the DEADc or HELICc domains alone (Fig 4K). Overall, these results indicate that DDX60 binds to TRAF6 mRNA via its HELICc domain; however, both the DEADc and HELICc domains are required for effective DDX60-driven expression of TRAF6 proteins.

To investigate the role of DDX60 in terms of association of transcripts and YTHDF1, we first investigated the binding of YTHDF1 to TRAF6 mRNA in the absence or presence of DDX60. As shown in Fig 5A, DDX60 depletion abrogated the interaction between YTHDF1 and TRAF6 mRNA. Further, we transfected YTHDF1-containing plasmid into IPEC-J2 cells either depleted of YTHDF1 only or both YTHDF1 and DDX60. We found that YTHDF1 could not rescue TRAF6 expression and p65 activation in YTHDF1-depleted cells in the absence of DDX60 (Fig 5B). These results indicated that the function of YTHDF1 in TRAF6 synthesis depends on DDX60. In addition, similar results from the TRAF6 and NF-κB luciferase reporter assay confirmed this conclusion (Fig 5C). Mechanistically, we employed constructed plasmids expressing various truncated fragments of YTHDF1 and DDX60, which revealed that YTHDF1–DDX60 interaction required the P/N/Q rich domain of YTHDF1 and the DEAD domain of DDX60 (Fig 5D and E). Thus, we conclude that the P/Q/N-rich domain of YTHDF1 and the DEAD domain of DDX60 are responsible for YTHDF1–DDX60 interaction and the function of YTHDF1 in TRAF6 synthesis.

**Fig 5.**
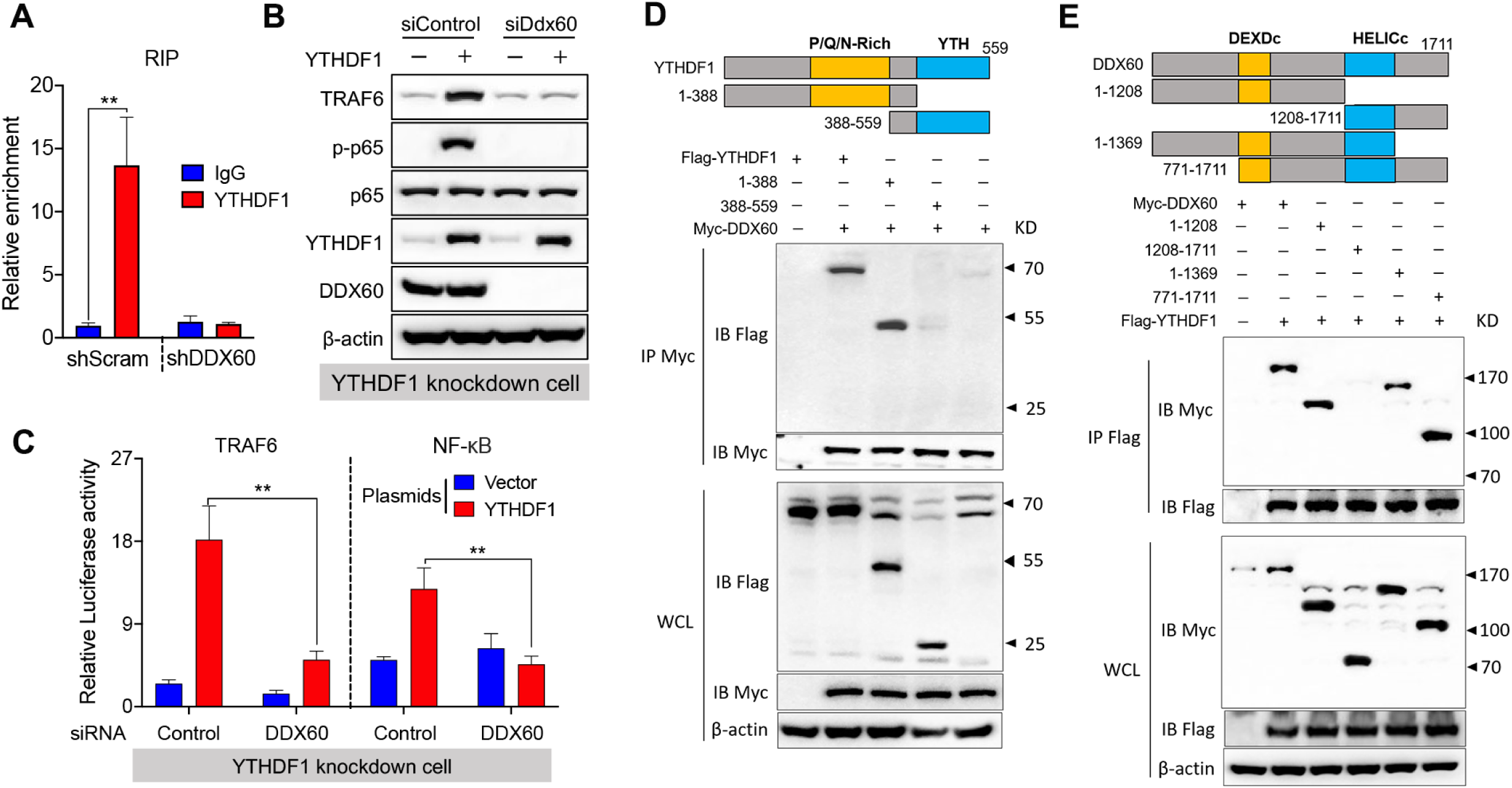
Interaction with DDX60 was critical for the role of YTHDF1 in TRAF6 protein synthesis. (A) RIP-qPCR analysis of the interaction between YTHDF1 and TRAF6 transcripts in IPEC-J2 cells with or without DDX60. (B) Rescue assay of YTHDF1 plasmid in LPS-treated YTHDF1 knockdown IPEC-J2 cells in the presence or absence of DDX60. (C) Luciferase activity of TRAF6 and NF-κB after co-transfection with the indicated plasmids in YTHDF1 knockdown IPEC-J2 cells subjected to DDX60 siRNA treatment. Luciferase activity is presented relative to Renilla luciferase activity. The data are expressed as the mean ± s.e.m.; ^**^*P* < 0.01, n = 3 biological replicates. (D-E) FLAG-tagged YTHDF1 partial fragments and/or Myc-tagged DDX60 expression vectors (D) or Myc-tagged DDX60 partial fragments and/or FLAG-tagged YTHDF1 expression vectors (E) were transfected into IPEC-J2 cells. After LPS stimulation for 6 h, immunoprecipitation was performed with Flag or Myc antibodies.

### YTHDF1 deficiency impedes the intestinal immune response *in vivo*

To further elucidate the function of YTHDF1 during the bacterial immune response, we created a targeted deletion of YTHDF1 in mice using the CRISPR/Cas9 system (Fig EV10A). Immunoblotting showed that the expression of YTHDF1 was abrogated in the jejunum of YTHDF1^−/−^ mice following infection (Fig EV10B). YTHDF1^−/−^ and YTHDF^+/+^ (i.e., WT) mice with similar body weights were selected and challenged with ETEC infection via oral gavage. Compared with YTHDF^+/+^ mice, YTHDF1^−/−^ mice exhibited considerably less weight loss (Fig 6A), which was also accompanied by a higher survival rate (Fig 6B) and longer colon length (Fig 6C). We next evaluated production of cytokines and found that ETEC infection induced increased production of inflammatory cytokines in YTHDF^+/+^ mice, which was largely compromised in YTHDF1^−/−^ mice (Fig 6D). Histological analysis showed a suppressed inflammatory response during ETEC infection accompanied by lower clinical disease scores in YTHDF1^−/−^ mice as compared with YTHDF^+/+^ mice (Fig 6E). Intestinal immune response is often accompanied by infiltration of inflammatory cells. As shown in Fig 6F, compared with YTHDF^+/+^ mice, YTHDF1^−/−^ mice exhibited reduced infiltration of immune cells (CD11b^+^, CD11c^+^, and F4/80^+^) in the jejunum upon ETEC infection. This finding was further supported by immunohistochemistry results in the colon (Fig EV10C). Finally, we evaluated TRAF6 synthesis *in vivo* and found that the TRAF6 protein level in the jejunum and colon from YTHDF1^−/−^ mice was significantly lower than the corresponding levels in YTHDF^+/+^ mice, albeit a similar level of TRAF6 mRNA was present (Fig 6G), suggesting that YTHDF1 plays a significant role in TRAF6 synthesis *in vivo*. These data convincingly indicate that YTHDF1 is vital to homeostatically sustain host antibacterial immunity.

**Fig 6.**
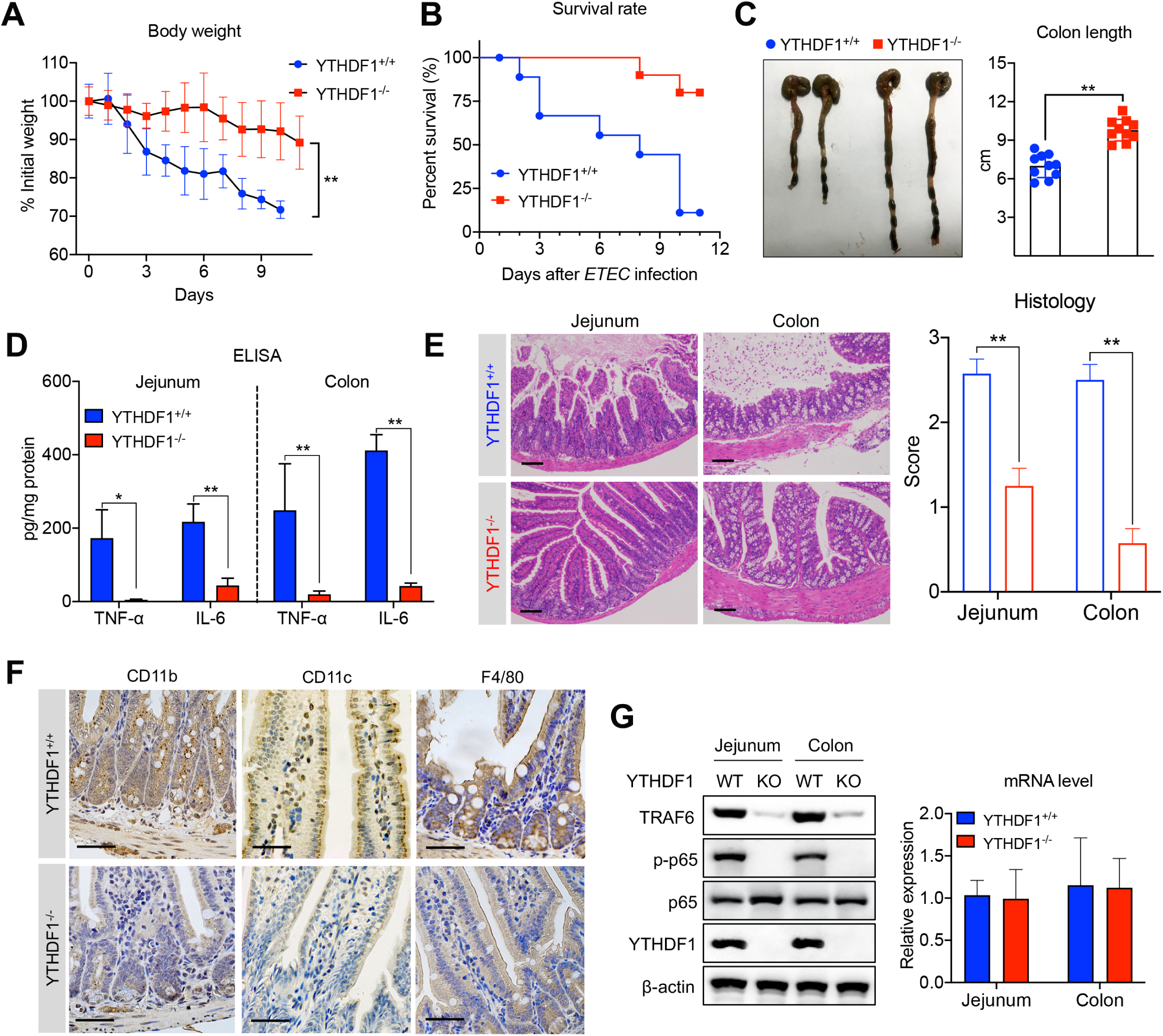
YTHDF1 deficiency reduced the bacterial immune response in the intestine *in vivo*. (A) Quantification of weight loss in YTHDF1^+/+^ and YTHDF1^−/−^ littermates after *ETEC* infection; n = 10 biological replicates for each genotype; \*\**P* < 0.01. (B) Survival curve of YTHDF1^+/+^ and YTHDF1^−/−^ mice following *ETEC* infection; n = 10 biological replicates for both YTHDF1^+/+^ and YTHDF1^−/−^ mice. (C) Representative gross images of colon from YTHDF1^+/+^ and YTHDF1^−/−^ littermates 3 days post-*ETEC* infection. (D) Cytokines in jejunum and colon of YTHDF1^+/+^ and YTHDF1^−/−^ mice were determined by ELISA. (E) Histological images of jejunum and colon from YTHDF1^+/+^ and YTHDF1^−/−^ mice after *ETEC* infection; n = 6 biological replicates for each genotype; scale bar: 50 mm. (F) Immunohistochemistry (IHC) showed infiltration of immune cells *in situ* in jejunum segments from *ETEC*-challenged YTHDF1^+/+^ and YTHDF1^−/−^ mice (brown staining, scale bar = 50 μm). (G) Results from the immunoblot analysis of total YTHDF1, TRAF6, p65 and p-p65 in the jejunum and colon from YTHDF1^+/+^ or YTHDF1^−/−^ mice. The right panel shows the mRNA level of TRAF6 in the same sample as indicated on the left. Results are presented in relation to those of GAPDH. The data are expressed as the mean ± s.e.m.; ^**^*P* < 0.01; n = 6 biological replicates.

## Discussion

Epigenetic factors have provided new insight into the pathogenesis of immune responses. Among many epigenetic modifications, RNA m^6^A modification is one of the research focuses, which have shown to be related to innate immunity (Goubau et al., 2015, Hao, Hao et al., 2019, Li, Tong et al., 2017, Shi et al., 2017). However, the specific function of m^6^A “readers” within the control of IEC immune responses and whether m^6^A is involved in these events remain unclear. Bacteria activate the immune response of IECs by interacting with LPS, which is capable of passing through the intestinal barrier. In this study, we established models with LPS and *ETEC* mimicking the immune response in IECs following bacterial infection. We showed that YTHDF1 regulated IEC immune response by directing TRAF6 translation in an m^6^A dependent manner. In addition, we identified DDX60 as a coregulator that facilitates YTHDF1 to recognize m^6^A on TRAF6 mRNA.

The biological function of m^6^A is mediated by its “readers”. Previous study has shown that infection of human foreskin fibroblasts by cytomegalovirus (HCMV) upregulates the level of YTHDF2 (Winkler, Gillis et al., 2019). We have shown that YTHDF1 was much more sensitive to *ETEC* infection than YTHDF2 or YTHDF3. Our data showed that depletion of YTHDF1 led to a suppressed intestinal immune response by blocking the NF-κB signaling pathway both *in vivo* and *in vitro*.

YTHDF1 was widely thought to enhance translation of its targets by interacting with initiation factors and facilitating ribosome loading (Wang et al., 2015). As revealed by Ribo-Seq data, the translation efficiency of m^6^A-modified transcripts was notably decreased in YTHDF1 knockdown IPEC-J2 cells in contrast to scramble control. We focused on the NF-κB signaling pathway, and observed that TRAF6 translation efficiency was significantly decreased, but not IκBα mRNA. Therefore, we speculated that TRAF6 mRNA was the target of YTHDF1 to govern the immune responses. However, a previous study showed that HCMV or dsDNA could trigger m^6^A-modified IFNβ production via the m^6^A methyltransferase METTL3 and demethylase ALKBH5 (Rubio, Depledge et al., 2018). These results indicated that YTHDF1 could utilize two different mechanisms to regulate the IEC immune response. First, YTHDF1 regulated the translation of TRAF6 mRNA to control the activation of NF-κB signaling pathway. Second, YTHDF1 initiated the translation of m^6^A-modified effector molecule mRNAs, such as IFNβ, to regulate their production directly.

To test whether m^6^A is involved in the IEC immune response, we measured methylation levels in LPS-stimulated cells with or without YTHDF1. Interestingly, following YTHDF1 depletion no increase in methylation was observed, likely because YTHDF1 is an m^6^A “reader” and not a methyltransferase. Decreased METTL3 and increased FTO levels in YTHDF1-knockdown cells suggested a feedback mechanism between YTHDF1 and m^6^A levels, which also substantiated our hypothesis. We further confirmed and characterized the m^6^A sites on TRAF6 mRNA by m^6^A-Seq. Consistent with the classical distribution of m^6^A on the gene structure (Dominissini, Moshitch-Moshkovitz et al., 2012), methylation generating m^6^A modifications on TRAF6 mRNA was concentrated in the vicinity of the stop codon at the conserved motif “RRACU”. To clearly illustrate the mechanism by which YTHDF1-m^6^A participates in the IEC immune response, we constructed TRAF6 plasmids either with or without the 3**′**UTR region (designated CDS-3**′**UTR or CDS, respectively). To our surprise, both the TRAF6 CDS-3**′**UTR and CDS-only constructs successfully overexpressed TRAF6 protein. These results indicated that m^6^A within the 3**′**UTR is not critically involved in YTHDF1-TRAF6 mRNA regulation. A similar finding was observed in TGF-β-induced SNAI1 expression, in which an m^6^A in the 3**′**UTR had no effect on its protein synthesis (Lin, Chai et al., 2019). Mao et al. found that removing CDS m^6^A from transcripts resulted in a further decrease in translation (Mao, Dong et al., 2019). Therefore, these results improved the previous supposition that m^6^A at the 3**′**UTR region mediated the regulatory role of YTHDF1 during translation (Wang et al., 2015, Zhang, Zhao et al., 2017). Taking advantage of variations of TRAF6 plasmids containing m^6^A site mutations, we found that the m^6^A modification was essential for TRAF6 protein synthesis but was not involved in its transcription. Considering the findings of the current study as well as the interaction between YTHDF1 and the EIF3 complex (Wang et al., 2015), we believe that a mediated loop structure formation is likely the key step for YTHDF1-m^6^A participation in translation, regardless of whether m^6^A is within the 3**′**UTR or the CDS near the stop codon.

Structural and binding studies have suggested that the YTH domain is a specific m^6^A “reader” (Xu, Wang et al., 2014, Zhu, Roundtree et al., 2014). We also found that YTHDF1 functioned by directly binding to TRAF6 transcripts at its YTH domain. However, our results suggest that loss of the P/Q/N rich domain failed to rescue YTHDF1 function in cytokine expression within YTHDF1-silenced IPEC-J2 cells, indicating that the intact YTHDF1 protein structure was required for TRAF6 protein synthesis. YTHDF family proteins share similar domain structures; however, in endometrial cancer cells YTHDF1 and YTHDF2 accurately target unique transcripts and regulate mRNA translation and decay, respectively (Shi et al., 2019). It is unknown as to how YTHDF1 or YTHDF2 recognize their specific m^6^A-marked targets accurately with the same domain. We hypothesized that a coregulator interacts with the P/Q/N-rich domain to ensure that each YTHDF1–TRAF6 transcript fulfills its specific role.

Based on the above hypothesis, we purified YTHDF1-interacted protein and identified the top candidates DDX60 as a coregulator in the YTHDF1-TRAF6 transcript interaction. Members of the DDX family of helicases broadly participate in the innate immune response as coregulators. It has been shown that DDX46 proteins bound to specific antiviral transcripts to negatively regulate the innate antiviral response (Zheng, Hou et al., 2017). As expected, the relationship among YTHDF1, DDX60, and TRAF6 was further confirmed through a YTHDF1 rescue assay in DDX60 and YTHDF1 double knockdown IPEC-J2 cells. Mechanistically, our data suggested that DDX60 binds to TRAF6 transcripts via its HELICc domain and interacts with the P/Q/N-rich domain of YTHDF1 through its DEAD helicase domain, which partially explains why YTHDF1 specifically recognizes its unique targets using the same domain as other YTHDF proteins. However, consequently, decreased mRNA level of TRAF6 was also observed in DDX60-depleted cells. We speculated that DDX60 recognizes mRNA in the nucleus and functions as an RNA carrier, which awaits further validation.

In summary, we identified a specific function and underlying mechanism for RNA m^6^A modification in regulating the intestinal immune response via the complex of YTHDF1, DDX60, and TRAF6 mRNA (Fig 7). This work not only provides novel insights into the molecular mechanisms underlying IEC immune response but also suggests novel strategies to cope with bacterial infections in the intestine.

**Fig 7.**
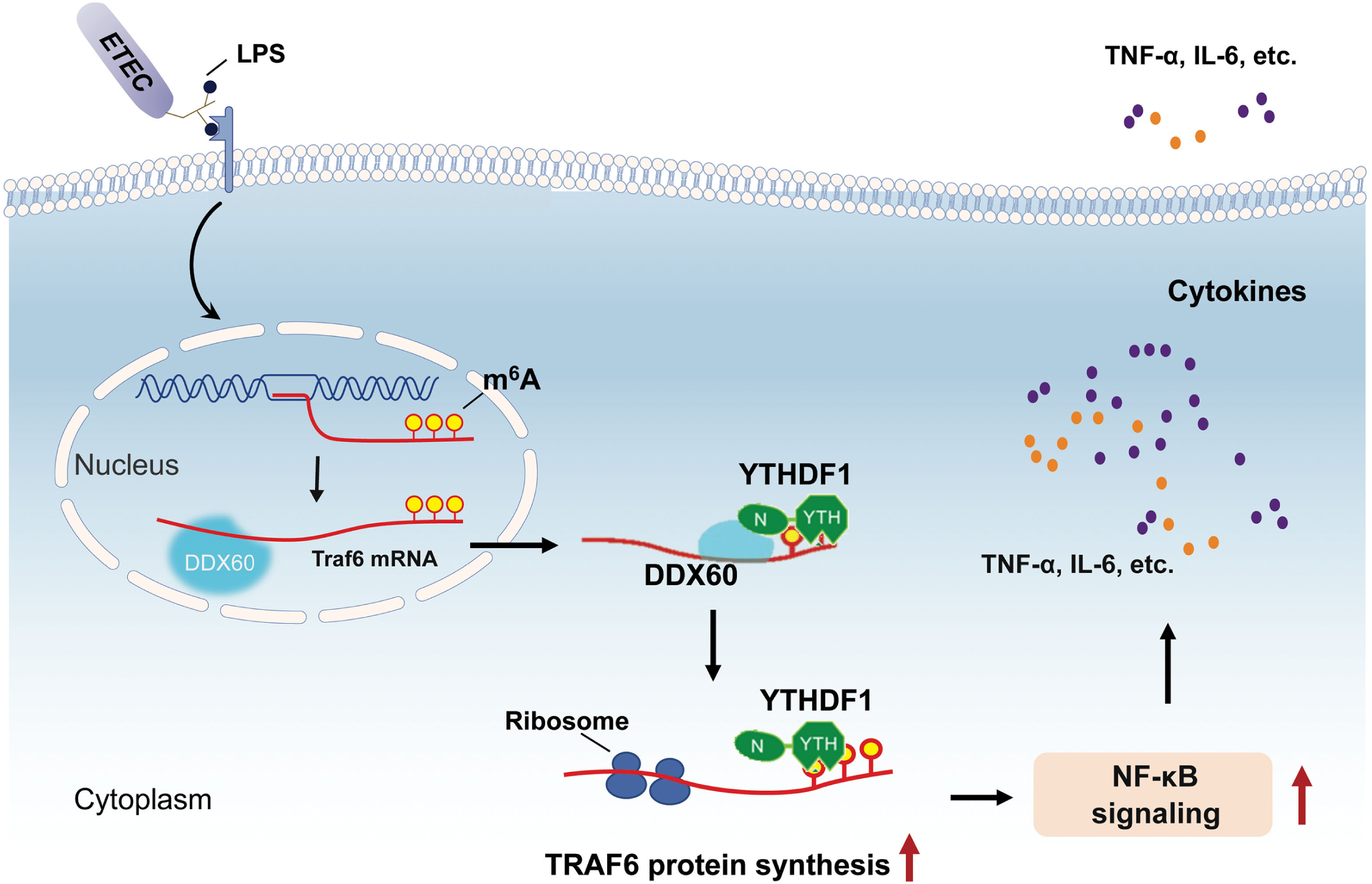
A working model in which YTHDF1 triggers the intestinal immune response against bacterial infection. DDX60 recruits YTHDF1 to recognize and positively regulate the translation of TRAF6 transcripts to drive intestinal immune responses against bacterial infection.

## Materials and Methods

### Cell culture

The porcine intestinal epithelial cell line IPEC-J2, human intestinal epithelial cell line Caco-2, and HEK 293T cells were purchased from the Cell Bank of the Chinese Academy of Sciences (Shanghai, China) and were cultured as described previously (Jiang, Sun et al., 2019, Zong, Zhao et al., 2019).

### Pathogens

*Escherichia coli* strain O111:B4 LPS (Sigma-Aldrich) and enterotoxigenic *Escherichia coli* K88 (ETEC) were provided by Dr. W. Fang (Zhejiang University, China) and cultured in Luria–Bertani (LB) broth (Aoboxing, China) or on LB agar plates.

### Reagents

Antibodies to the following were used in this study: anti-m^6^A (Synaptic Systems, 202003), YTHDF1 (Proteintech, 66745-1-lg), TRAF6 (abcam, ab181622), β-actin (HuaAn, R1207-1), anti-Flag (Sigma-Aldrich, F3165), anti-Myc (abcam, ab32), DDX60 (abcam, ab139807), MyD88 (Proteintech, 23230-1-AP), phosphor-p65 (Cell Signaling Technology, 3033), phosphor-IκBα (Cell Signaling Technology, 2859), TRIF (Proteintech, 23288-1-AP), p65 (Proteintech, 10745-1-AP), IκBα (Proteintech, 10268-1-AP), TRAF3 (Proteintech, 18099-1-AP), phospho-IKKα/β (Cell Signaling Technology, 2078), IKKα (HuaAn, ER30911), IKKβ (HuaAn, ER1706-13), p38 (HuaAn, ET1602-26), phospho-p38 (Cell Signaling Technology, 9211), phospho-JNK (HuaAn, RT1488), JNK (HuaAn, RT1550), ERK1/2 (Cell Signaling Technology, 9102), phospho-ERK1/2 (HuaAn, ET1610-13), normal rabbit IgG (Cell Signaling Technology, 2729), and normal mouse IgG (Cell Signaling Technology, 5946).

### Mice and treatments

All animal procedures were performed in accordance with the Guide for the Care and Use of Laboratory Animals in Zhejiang University. (Hangzhou, China). C57BL6/J mice (6–8 weeks old) were obtained from the Laboratory Animal Center of the Chinese Academy of Sciences (Shanghai, China). YTHDF1-deficient (YTHDF1^−/−^) mice on a C57BL/6J background were generated using CRISPR-Cas9 system, which were obtained from Bin Shen Laboratory in Nanjing Medical University (Nanjing, China). The oligonucleotides for sgRNAs were as follows: 1, TACCTGTCCAGTTACTATCC; 2, GGCACCATGGTCCACTGGAG. YTHDF1^+/+^ and YTHDF1^−/−^ mice were infected with 100 μl of the prepared bacterial suspension at a dose equivalent to 10^9^ CFU/mouse by oral gavage. Total experimental period was 11 days. C57BL/6 mice were sacrificed at day 3 post-infection.

### Lentiviral short hairpin RNAs, siRNA and transfection

YTHDF1- and DDX60-knockdown IPEC-J2 cells were generated by using lentiviral short hairpin (sh) RNAs (shRNAs); shRNA targeting sequences and siRNA are listed in Table EV1. Plasmids or siRNA were transfected into cells with Lipofectamine 2000 (Invitrogen) according to the manufacturer’s instructions.

### Molecular cloning of related genes

Related genomic cDNA were cloned from total RNAs extracted from IPEC-J2 cells by RT-PCR. FLAG-tagged YTHDF1 was cloned into pCMV-Tag 4 vector (Agilent, #211174); Myc-tagged DDX60 was cloned into pRK-5 vector (BD PharMingen, 556104). We generated truncation plasmids based on full-length proteins. All other plasmids, including full-length, coding sequence, and mutation plasmids were generated by GenScript, which were then cloned into FLAG-pcDNA3.1 vector. All plasmids were confirmed by DNA sequencing. The corresponding primers used in this study are listed in Table EV1.

### RNA extraction, cDNA synthesis, and qPCR

Total RNA was isolated using TRIzol reagent (Invitrogen), and poly(A)^+^ mRNA was enriched from total RNA with GenElute mRNA Miniprep Kit (Sigma-Aldrich). The cDNA was reverse transcribed using M-MLV reverse transcriptase (Thermo Fisher Scientific) with 2 μg of RNA from each sample. Next, quantitative PCR (qPCR) was performed as previously described (Zong et al., 2019). GAPDH or β-actin was used as endogenous control, and each reaction was run in triplicates. Gene copy numbers were then calculated using the following formula: number of copies = (DNA quantity × 6.022 × 10^23^)/(DNA length × 1 × 10^9^ × 650). The primer sequences used for qPCR are listed in Supplementary Table EV1.

### RNA immunoprecipitation (RIP) assay

RIP was performed according to the native RIP protocol as previously described (Gagliardi & Matarazzo, 2016, Shen, Zhang et al., 2018). Briefly, cell lysates were prepared in polysome lysis buffer (100 mM KCl, 5 mM MgCl2, 10 mM HEPES pH 7.0, and 0.5% NP40) supplemented with DTT, protease inhibitor cocktail (APExBIO), and RNase inhibitor (Promega) on ice. Cell lysates were incubated with antibody and protein A/G magnetic beads (Sigma) from 3 h to overnight at 4°C under rotation conditions. Ten percent of the cell lysate supernatant was saved for input analysis. Beads were washed four times and the RNA released with proteinase K treatment for 30 min at 55°C. Precipitated RNA was extracted and analyzed by qPCR. The primer sequences used are listed in Table EV1.

### m^6^A dot blot

The m^6^A dot blots were conducted as previously described (Zhou, Wan et al., 2015, Zong et al., 2019). Equal amounts of poly(A)^+^ mRNA was spotted on an Amersham Hybond-N+ membrane (GE Healthcare), followed by UV crosslinking. After blocking, the membrane was incubated with anti-m^6^A antibody overnight at 4°C. The dots were visualized using a ChemiScope series 3400 Mini Imaging System. Methylene blue (Sigma-Aldrich) staining was used to show the amount of total RNA on the membrane.

### Immunoblot

Cells were lysed on ice in SDS-PAGE sample buffer [50 mM Tris (pH 6.8), 100 mM DTT, 2% SDS, 0.1% bromophenol blue, and 10% glycerol]. Proteins were transferred onto a polyvinylidene fluoride membrane (MilliporeSigma, Billerica, MA) and were blotted and quantified as described previously (Zong et al., 2019).

### Luciferase reporter assays

IPEC-J2 cells were plated in 24-well plates and then were transfected with pGL4.21 firefly luciferase reporter plasmid and Renilla luciferase plasmid, together with various amounts of the appropriate control or protein-expressing plasmid(s). The luciferase activity was measured using the Dual-Luciferase Reporter Assay System (Promega). Reporter gene activity was determined by normalization of firefly luciferase activity to Renilla luciferase activity.

### Protein co-immunoprecipitation

The cell pellet was resuspended with 2 volumes of IP lysis buffer (Pierce) supplemented with cocktail protease inhibitor (APExBIO), incubated on ice for 10 min, and then incubated with primary antibody and Protein A/G magnetic beads at 4°C overnight. The washed beads were resuspended in SDS sample buffer and analyzed by immunoblotting or tandem mass spectrometry analysis.

### Tandem mass spectrometry analysis

Peptide samples were analyzed using an Orbitrap Elite hybrid mass spectrometer (Thermo Fisher) by Novogene (Beijing, China). Briefly, the purified protein samples were separated by SDS-PAGE and visualized by silver staining. The proteins were reduced, alkylated, and digested as previously described (Shevchenko, Tomas et al., 2006). The extracted samples were dried, resuspended in 0.1% trifluoroacetic acid, desalted with C18 ZipTips, dried again, and dissolved in 0.1% formic acid. Samples were separated on a C18 analytical column using a two-solvent system.

### Cytokine multiplex assay

Porcine cytokine antibody arrays (QAP-CYT-1) were obtained from and subsequently conducted by RayBiotech (Guangzhou, China). Serum-free media from IPEC-J2 cultures or equivalent amounts of mouse intestinal tissue homogenate were used. Intensity was normalized to internal positive controls for comparison.

### Cell treatments and ELISA

The cells were cultured in six-well dishes until they reached approximately 80% confluence and then treated with LPS (50 μg/ml) or *Escherichia coli* K88 (MOI 10:1) for varying durations. Untreated cells were used as controls. The concentration of cytokines released into the supernatant or in intestines was measured via specific IL-6 or TNF-α ELISA (Quantikine ELISA Kit; Proteintech).

### Histology and immunohistochemistry

Excised intestine from euthanized mice was fixed in 4% paraformaldehyde overnight at 4°C. Samples were then sent to Servicebio (Wuhan, China) for histological and immunohistochemical analysis. Briefly, samples embedded in paraffin were deparaffinized and processed through a graded series of alcohol. Then, the sections were incubated with primary antibody followed by peroxidase-conjugated secondary antibody. Sections were scored by a blinded observer based on goblet cell depletion, leukocyte infiltration, and submucosal inflammation on a point scale of 0–3, with 0 representing no pathology and 3 indicating the most severe pathology.

### Ribosome-protected fraction isolation, Ribo-Seq and analysis

The ribosome-protected fraction isolation, library construction, high-throughput sequencing and analysis were performed by Novogene (Beijing, China). In brief, ribosome-protected mRNA was obtained by MNase digestion and RNA purification (Ingolia, Brar et al., 2012, Reid, Shenolikar et al., 2015). In total, 3 μg of ribosome-protected mRNA was used to prepare a library with the NEBNext Small RNA Library Prep Set (New England Biolabs) according to the manufacturer’s instructions. After PCR amplification, the samples were used for quality control and deep sequencing with an Illumina HiSeq 2000.

### m^6^A sequencing and analysis

Library preparation of m^6^A Sequencing and high-throughput sequencing were performed by Novogene (Beijing, China). The m^6^A immunoprecipitation and preparation of libraries were performed as previously described (Dominissini et al., 2012). Purified RNA fragments were used for library construction with the NEBNext Ultra RNA Library Prep Kit and were sequenced with Illumina HiSeq 2000. Sequencing reads were aligned to the Sscrofa11.1 by BWA, and the m^6^A peak calling was performed by MACS2.

### RNA sequencing and analysis

Library preparation of RNA sequencing and high-throughput sequencing were performed by Novogene (Beijing, China). Raw data were filtered, corrected, and mapped to locust genome sequences using the HISAT software. Gene expression levels were measured using the criteria of reads per kb per million mapped reads. DEGs were detected using the DESeq software (Trapnell, Williams et al., 2010). Genes with an adjusted *P*-value of <0.05 were described as being differentially expressed. Kyoto Encyclopedia of Genes and Genomes (KEGG) enrichment was performed by the Clusterfile software. Significance analysis was performed by Fisher’s Exact Test.

### Statistical analysis

Results were expressed as means ± s.e.m. For multiple groups, one-way ANOVA was used to determine statistical differences, and t-tests were employed between two groups; ^*^*P* < 0.05, ^**^*P* < 0.01. Data were analyzed with GraphPad Prism Software. Each experiment was performed at least in triplicates unless otherwise stated.

## Acknowledgements

The authors thank the Electronic Microscopy Center and the Agricultural, Biological, and Environmental Test Center of Zhejiang University for assistance with the confocal microscopy work. The authors thank Shu Zhu’s lab in Zhejiang University for assistance with the *ETEC* infection. This work was supported by the National Natural Science Foundation of China (Grants 31630075).

## Author contributions

X.Z and Y.Z.W designed the research; X.X, Q.J, H.W performed experiments and collected data; X.Z wrote the manuscript; B.S and F.Q.W supported technical; Z.Q.L and M.L.J reviewed the manuscript.

